# Changes in brain morphology and working memory capacity over childhood

**DOI:** 10.1101/069617

**Authors:** Joe Bathelt, Susan Gathercole, Amy Johnson, Duncan Astle

## Abstract

Developmental improvements in working memory are important in the acquisition of new skills, like reading and maths. Current accounts of the brain systems supporting working memory rarely take development into account. However, understanding the development of these skills, and in turn where this development can go awry, will require more sophsiticated neuropsychological accounts that fully consider the role of development. The current study investigated how structural brain correlates of components of the working memory system change over developmental time. Verbal and visuospatial short-term and working memory were assessed in 153 children between 6 and 16 years and latent components of the working memory system were derived using principal component analysis. Further, fractional anisotropy and cortical thickness maps were derived from T1-weighted and diffusion-weighted MRI and processed using eigenanatomy decomposition, an advanced dimensionality reduction method for neuroimaging data. We were then able to explore how the structural brain correlates of working memory gradually shifted across childhood. Regression modelling indicated greater involvement of the corpus callosum and posterior temporal white matter in younger children for performance associated with the executive part of the working memory system, while thickness of the occipitotemporal cortex was more predictive in older children. These findings are consistent with an account in which increasing specialisation leads to shifts in the contribution of neural substrates over developmental time, from early reliance on a distributed system supported by long-range connections to later reliance on specialised local circuitry. Furthemore, our findings emphasise the importance of taking development into account when considering the neural systems that support complex cognitive skills, like working memory.

## 2 Introduction

Working memory is a limited-capacity system for retaining and processing information over brief periods of time. It plays an important role in the acquisition of complex cognitive skills (Cowan, 2013) such as reading (Cain, Oakhill, & Bryant, 2004), maths (Dumontheil & Klingberg, 2011), and other school subjects (Gathercole, Pickering, Knight, & Stegmann, 2003, Clair-Thompson & Gathercole, 2006). Deficits in working memory have been identified across a range of neurodevelopmental disorders, including attention deficit hyperactivity disorder (Martinussen, Hayden, Hogg-Johnson, & Tannock, 2005, Holmes et al., 2014), dyslexia (Smith-Spark & Fisk, 2007), dyscalculia (Rotzer et al., 2009, Szucs et al., 2013), and language disorders (Gathercole & Baddeley 1989; Archibald & Gathercole 2006; Montgomery, 2000, Ellis Weismer, Evans, & Hesketh, 1999).

Working memory develops gradually through early and middle childhood (Huizinga, Dolan, & Molen, 2006, Gathercole, Pickering, Ambridge, & Wearing, 2004, Siegel & Ryan, 1988). It is assumed that this development reflects the maturation of the brain system supporting this skill in adulthood. However, understanding the potential causes of working memory impairments in childhood necessitates a neuropsychological account that incorporates developmental change. Currently we have no detailed understanding of how age-related changes in brain organisation support developmental improvements in working memory. The purpose of this study is to redress this.

### 2.1 Working Memory and its development

There are many theoretical accounts of working memory. The influential multicomponent model of working memory advanced by Baddeley and Hitch (A. D. Baddeley & Hitch, 1974) consists of three subcomponents: two domain-specific stores and a central executive. The stores are specialized for the retention of material in either phonological (A. Baddeley, 1987) or visuo-spatial format (A. D. Baddeley & Lieberman, 1980, Logie, 1986). The central executive is a system responsible for a range of regulatory functions, including attention, the control of action, and problem solving (A. Baddeley, 1996).

There have been many refinements of the original model (A. Baddeley, 2000, 2003, 2012; Burgess & Hitch, 1996), and several new accounts. Some of these elaborate on specific mechanisms within working memory. For instance, Engle and colleagues add inhibitory processes that protect activated memory traces from disruption (Engle, 2002, Kane, Conway, Hambrick, & Engle, 2007). Other models integrate short-term memory with long-term memory, suggesting that working memory represents long-term memory in an activated state (Cowan, 1988, 1999; Oberauer 2002), and activation is guided by an attentional mechanism. Others have extended the scope of WM to encompass other processes that include updating (Ecker et al., 2010; Schmiedek et al., 2014; Shelton et al, 2010), set shifting and relational binding (Oberauer et al., 2003; von Bastian & Oberauer, 2013) and fluid intelligence (Engle et al., 1999). In short, there exists a rich literature in which the specific cognitive mechanisms underlying working memory in adulthood are keenly debated.

Multiple tasks are needed to establish the underlying latent factor structure supporting working memory performance rather than simply characterise the cognitive structure of individual tasks. Using this individual differences approach, the three-factor structure has been robustly reproduced across multiple studies and age groups (T. P. Alloway, Gathercole, Willis, & Adams, 2004; Kane et al., 2004; Bayliss et al., 2003; Hornung, Brunner, Reuter, & Martin, 2011). In each case, the best-fitting model consisted of two domain-specific storage processes and an additional executive component similar to those outlined in the original Baddeley and Hitch (A. Baddeley, 2003, 2012) account. These components are already detectable in preschool-age children (T. P. Alloway et al., 2004) and their configuration remains stable throughout childhood (Gathercole et al., 2004). Despite this stable factor structure, overall working memory performance changes substantially over childhood (Huizinga et al., 2006, Gathercole et al., 2004, Siegel & Ryan, 1988), increasing linearly from 6 years until adult performance is reached in adolescence (Gathercole et al., 2004, Luciana, Conklin, Hooper, & Yarger, 2005). Different cognitive mechanisms may contribute to improvements across different periods (Huizinga et al., 2006, Gathercole et al., 2004, Siegel & Ryan, 1988). These may include increased storage capacity (Cowan, Ricker, Clark, Hinrichs, & Glass, 2014), improvements in attention (Barrouillet, Gavens, Vergauwe, Gaillard, & Camos, 2009, Tam, Jarrold, Baddeley, & Sabatos-DeVito, 2010), and changes in rehearsal strategy (Hitch, Halliday, Schaafstal, & Heffernan, 1991, Gathercole, Adams, & Hitch, 1994).

### 2.2 Neural correlates of working memory

The developmental period associated with increases in working memory is also accompanied by pronounced changes in brain structure. These include decreasing cortical thickness (Sowell, 2004) and increasing myelination of white matter tracts (Dean et al., 2014). Functional neuroimaging studies suggest that improvements in working memory are accompanied by some re-organisation in brain networks: In adults, a specialised network including bilateral parietal, cingulate, and prefrontal areas has been found to show increased blood oxygenation during working memory tasks (Owen, McMillan, Laird, & Bullmore, 2005, Wager & Smith, 2003). Children show activation in a similar set of regions (Thomason et al., 2009) and also in additional outside of the core processing network observed in adults (Vogan, Morgan, Powell, Smith, & Taylor, 2016, Ciesielski, Lesnik, Savoy, Grant, & Ahlfors, 2006).

Research on structural neural correlates of working memory is more limited, but where studies exist they broadly concur with the functional findings. Frontal and parietal grey matter volume (Mahone, Martin, Kates, Hay, & Horska, 2009, Rossi et al., 2013), and temporal and parietal connections of the corpus callosum (Treble et al., 2013), are significant predictors of a participant’s working memory capacity. However, these studies either investigate narrow age ranges or statistically correct for the effect of age. As a result, little is known about how structural brain changes support the development of particular cognitive skills like working memory. Furthermore, the majority of previous studies have used performance on individual tasks to measure working memory ability (see (Poldrack & Yarkoni, 2016) for a detailed discussion). This approach has two key limitations. First, it is widely accepted that multiple underlying components underpin performance (Conway, Cowan, Bunting, Therriault, & Minkoff, 2002, T. P. Alloway et al., 2004, Clair-Thompson & Gathercole, 2006, Oberauer, Süß, Schulze, Wilhelm, & Wittmann, 2000). Second, scores on individual tests also reflect task-specific components that may be unrelated to WM demands (such as proficiency in the stimulus domain from which the stimuli are drawn, Dark & Benbow, 1994) as well as significant levels of error. The purpose of the current study was to redress these two gaps in the literature by i) exploring how structural brain correlates of working memory, in terms of both grey and white matter, change over developmental time; and ii) using multiple behavioural assessments alongside a theory-driven factor analysis, to differentiate the neural correlates of robustly determined cognitive components of WM.

## 3 Methods

Our analysis approach used data reduction techniques to reduce raw behavioural and neuroimaging measures to underlying statistical components. We then explored how the underlying cognitive factors of the working memory system were associated with structural brain components, and the extent to which these relationships were moderated by developmental stage (i.e. age). A schematic summary of this approach can be seen in Figure 1. The computer code used for data processing and statistical analysis is available online (https://github.com/joebathelt/WorkingMemory_and_BrainStructure_Code).

**Figure 1:**
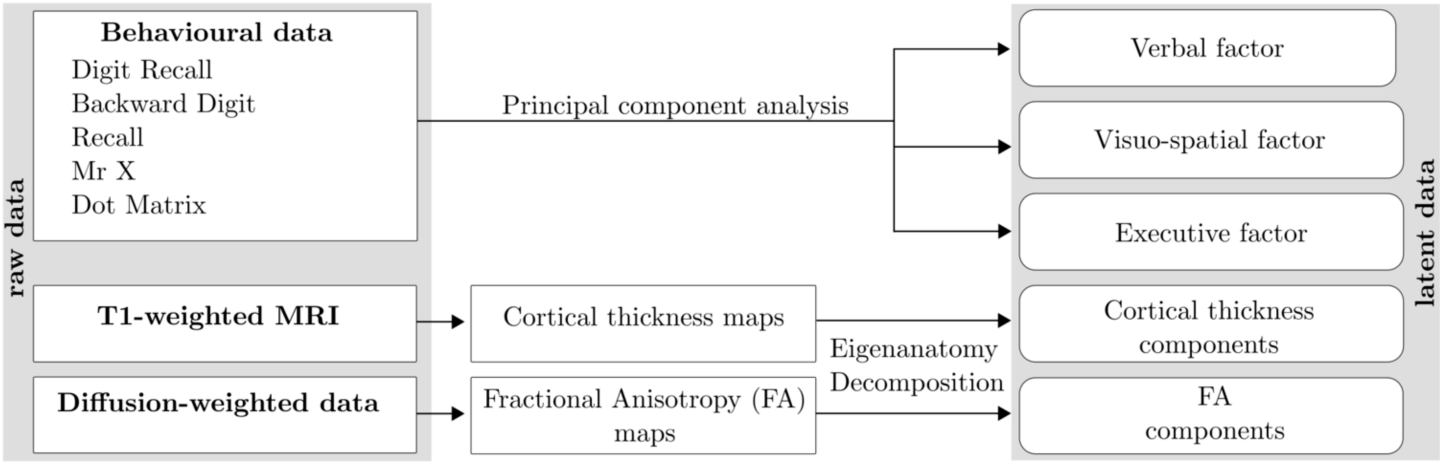
Overview of processing steps from raw to latent data. Raw behavioural data were decomposed with principal component analysis (PCA) to derive factor scores that corresponded to a verbal, visuo-spatial, and executive factor. Dimensionality reduction was also applied to cortical thickness maps and FA maps derived from T1-weighted and diffusion-weighted MRI data to obtain Eigenanatomy components.

### 3.1 Participants

The data for the current study were taken from two large-scale studies at the MRC Cognition and Brain Sciences Unit. These two studies had different recruitment criteria but when combined, provide a large sample of children with working memory scores whose distributional properties closely approximated the standardisation sample. The first study was the Centre for Attention, Learning, and Memory (CALM) research clinic. At the clinic, children aged between 5 and 18 years were recruited on the basis of ongoing problems in attention, learning and memory identified by professionals working in schools or specialist children’s community services. If parents expressed interest in participating in the research study, the professionals made an initial referral that was followed-up by CALM staff to discuss the nature of the child’s problems. If difficulties in one or more areas of attention, learning or memory were indicated by the referrer, the family were invited to the CALM clinic at the MRC Cognition and Brain Sciences Unit in Cambridge for an assessment that lasted approximately 3 hours. The assessment included the working memory battery reported here. Exclusion criteria for referrals were significant or severe known problems in vision or hearing that were uncorrected, and having a native language other than English. This study was approved by the local NHS research ethics committee (Reference: 13/EE/0157). Written parental consent was obtained and children provided verbal assent.

The second study investigated the neural, cognitive, and environmental markers of risk and resilience in children. Children between 7 and 12 years attending mainstream school in the UK with normal or corrected-to-normal vision or hearing and no history of brain injury were recruited via local schools and through advertisement in public places (childcare and community centres, libraries). Participating families were invited to the MRC Cognition and Brain Sciences Unit for a 2-hour assessment that included the working memory battery reported here. Participants received monetary compensation for taking part in the study. This study was approved by the Psychology Research Ethics Committee at the University of Cambridge (Reference: 2015.11). Parents provided written informed consent.

MRIs were obtained from 153 children between 6 and 16 years (96 boys, age in months: M=115.79, SD=23.779). 31 children were excluded from cortical thickness analysis because the T1-weighted data was not usable due to participant movement. 41 children were excluded from analysis of diffusion-weighted data due to head movement above 3mm in the DWI sequence. Residual movement estimates were included as a nuisance variable in regression models. As these measures did not influence the results, they were omitted from the reported models.

### 3.2 Working Memory Assessment

The Digit Recall, Backward Digit Recall, Dot Matrix, and Mr X task of the Automatic Working Memory Assessment (AWMA) (T. Alloway, 2007, T. P. Alloway, Gathercole, Kirkwood, & Elliott, 2008) were administered individually. In Digit Recall, children repeat sequences of single-digit numbers presented in an audio format. In Backward Digit Recall, children repeat the sequence in backward order. These tasks were selected to engage verbal short-term and working memory respectively. For the Dot Matrix task, the child is shown the position of a red dot for two seconds in a series of four by four matrices and has to recall this position by tapping the squares on the computer screen. In the Mr. X task, the child is shown a series of Mr. X figures and has to identify whether they are holding the ball in the same or different hands. One Mr. X is rotated in each trial. The child then has to recall the location of the ball in Mr. X’s hand by pointing to one of eight compass points. These tasks were aimed at tapping short-term and working visuo-spatial memory.

Standardised scores established that the sample performed at expected levels for their age (i.e. mean of 100 and a standard deviation of 15, Digit Recall: mean = 96.39; std = 16.32; Backward Digit Recall: mean = 94.61, std = 12.671; Dot Matrix: mean = 98.29, std = 15.595; Mr X: mean = 99.32, std = 15.69).

In order to reconstruct the latent variable structure of working memory from the assessment data, principal component analysis was applied. This was carried out using the ‘principal’ function of the psych package v1.5.1 (http://personality-project.org/r) in R v3.1.3 (R Development Core Team, 2008). Varimax rotation was used to create orthogonal factors (Kaiser, 1958). A 3-factor solution provided the best fit with theoretical predictions and explained a large proportion of variance in the assessment scores. Mahalanobis distance was computed to detect outliers in the assessment data, but no data point exceeded the recommended cut-off at 3 degrees of freedom.

### 3.3 MRI data acquisition

Magnetic resonance imaging data were acquired at the MRC Cognition and Brain Sciences Unit, Cambridge U.K. All scans were obtained on the Siemens 3 T Tim Trio system (Siemens Healthcare, Erlangen, Germany), using a 32-channel quadrature head coil. The imaging protocol consisted of two sequences: T1-weighted MRI and a diffusion-weighted sequence.

T1-weighted volume scans were acquired using a whole brain coverage 3D Magnetisation Prepared Rapid Acquisition Gradient Echo (MP RAGE) sequence acquired using 1mm isometric image resolution. Echo time was 2.98 ms, and repetition time was 2250 ms.

Diffusion scans were acquired using echo-planar diffusion-weighted images with an isotropic set of 60 non-collinear directions, using a weighting factor of b = 1000s*mm^−2^, interleaved with 4 T2-weighted (b = 0) volumes. Whole brain coverage was obtained with 60 contiguous axial slices and isometric image resolution of 2mm. Echo time was 90 ms and repetition time was 8400 ms.

### 3.4 Processing of diffusion-weighted data

Diffusion imaging makes it possible to quantify the rate of water diffusion in the brain. In the parallel bundles of white matter, diffusion is stronger along the fibre orientation, but is attenuated in the perpendicular direction. This can be summarized by the metric of fractional anisotropy (FA), which is a scalar value between 0 and 1 describing the degree of anisotropy of the diffusion at every voxel. Developmental studies show steady increases in FA between childhood and adulthood (Imperati et al., 2011, Muftuler et al., 2012, Westlye et al., 2009), which is likely to reflect increased myelination (Dean et al., 2014).

A number of processing steps are necessary to derive FA maps from diffusion-weighted volumes. In the current study, diffusion-weighted MRI scans were converted from the native DICOM to compressed NIfTI-1 format using the dcm2nii tool http://www.mccauslandcenter.sc.edu/mricro/mricron/dcm2nii.html. Subsequently, the images were submitted to the DiPy v0.8.0 implementation (Garyfallidis et al., 2014) of a non-local means de-noising algorithm (Coupe et al., 2008) to boost signal to noise ratio. Next, a brain mask of the b0 image was created using the brain extraction tool (BET) of the FMRIB Software Library (FSL) v5.0.8. Motion and eddy current correction were applied to the masked images using FSL routines. The corrected images were re-sliced to 1mm resolution with trilinear interpolation using in-house software based on NiBabel v2.0.0 functions (http://nipy.org/nibabel/). Finally, fractional anisotropy maps were created based on a diffusion tensor model fitted through the FSL dtifit algorithm (Behrens et al., 2003, Johansen-Berg et al., 2004).

For comparison across participants, we created a study-specific FA-templates based on all available images using Advanced Normalization Tools (ANTs) algorithms (Lawson, Duda, Avants, Wu, & Farah, 2013, B. B. Avants et al., 2014), which showed the highest accuracy in software comparisons (Klein et al., 2009, Murphy et al., 2011, Tustison et al., 2014). Individual images were transformed to template space using non-linear registration with symmetric diffeomorphic normalization as implemented in ANTs (B. Avants, Epstein, Grossman, & Gee, 2008). Next, the images were eroded twice with a 3mm sphere to remove brain edge artefacts using FSL maths.

### 3.5 Processing of T1-weighted data

Another measure of brain development that can be derived from neuroimaging data is cortical thickness (Giedd & Rapoport, 2010, Gogtay et al., 2004). Cortical thickness is defined as the distance between the outer edge of cortical grey matter and subcortical white matter (Fischl & Dale, 2000). To obtain thickness measure moments from anatomical MRI data, T1-weighted volumes were initially co-registered with MNI152 space using rigid co-registration in order to obtain good initial between-subject alignment and optimal field of view. Next, all images were visually inspected and images with pronounced motion artefact were removed from further analysis (n=31, 20.25% of the acquired data). The remaining data was submitted to the automatic ANTs cortical thickness pipeline (antsCorticalThickness). Details about the processing pipeline and thickness estimation are described in (Tustison et al., 2014) and (Das, Avants, Grossman, & Gee, 2009). Tissue priors were taken from the OASIS-TRT-20 template (http://www.mindboggle.info/data.html#mindboggle-software-data). Subsequently, images in template space were smoothed using a 10mm full width at half maximum (FWHM) Gaussian kernel and resampled to 2mm resolution. A thickness mask was created by averaging all images and binarizing the resulting mean image at a threshold of 0.1.

### 3.6 Eigenanatomy Decomposition

Traditional univariate approaches like voxel-based morphometry (VBM) fit a statistical model for every voxel in a brain image. Because of the large number of voxels in a typical imaging protocol, this approach necessitates correction for a very large number of comparisons (T1-volumes in the current study contained over 1 million voxels), resulting in a dramatic loss of statistical power. However, effects are typically spread over areas that are larger than 1 voxel. Multivariate approaches are better suited to reduce the dimensionality of the data to the information contained in the data itself before statistical comparisons are applied. Eigenanatomy Decomposition is a novel method for data-driven dimensionality reduction of neuroimaging data that adds sparseness and smoothness constraints for better anatomical interpretability in comparison to standard spatial principal component analysis (Kandel, Wang, Gee, & Avants, 2015). Cortical thickness masks and FA images were processed using the ANTsR v0.3.2 implementation of the Eigenanatomy Decomposition algorithm (Kandel et al., 2015). Parameters for Eigenanatomy Decomposition were adopted from published work, i.e. decomposition into 32 components with sparseness of 1/32 with 20 iterations, a L1 penalty with gradient step size 0.5, a smoothing kernel of 1 voxel, and a minimum cluster size of 1000 voxels for each eigenvector. For statistical analysis, the mean value of each brain morphology measure (FA, cortical thickness) within each eigenanatomy component was calculated. See Figure 2 for an illustration of the resulting parcellation.

**Figure 2:**
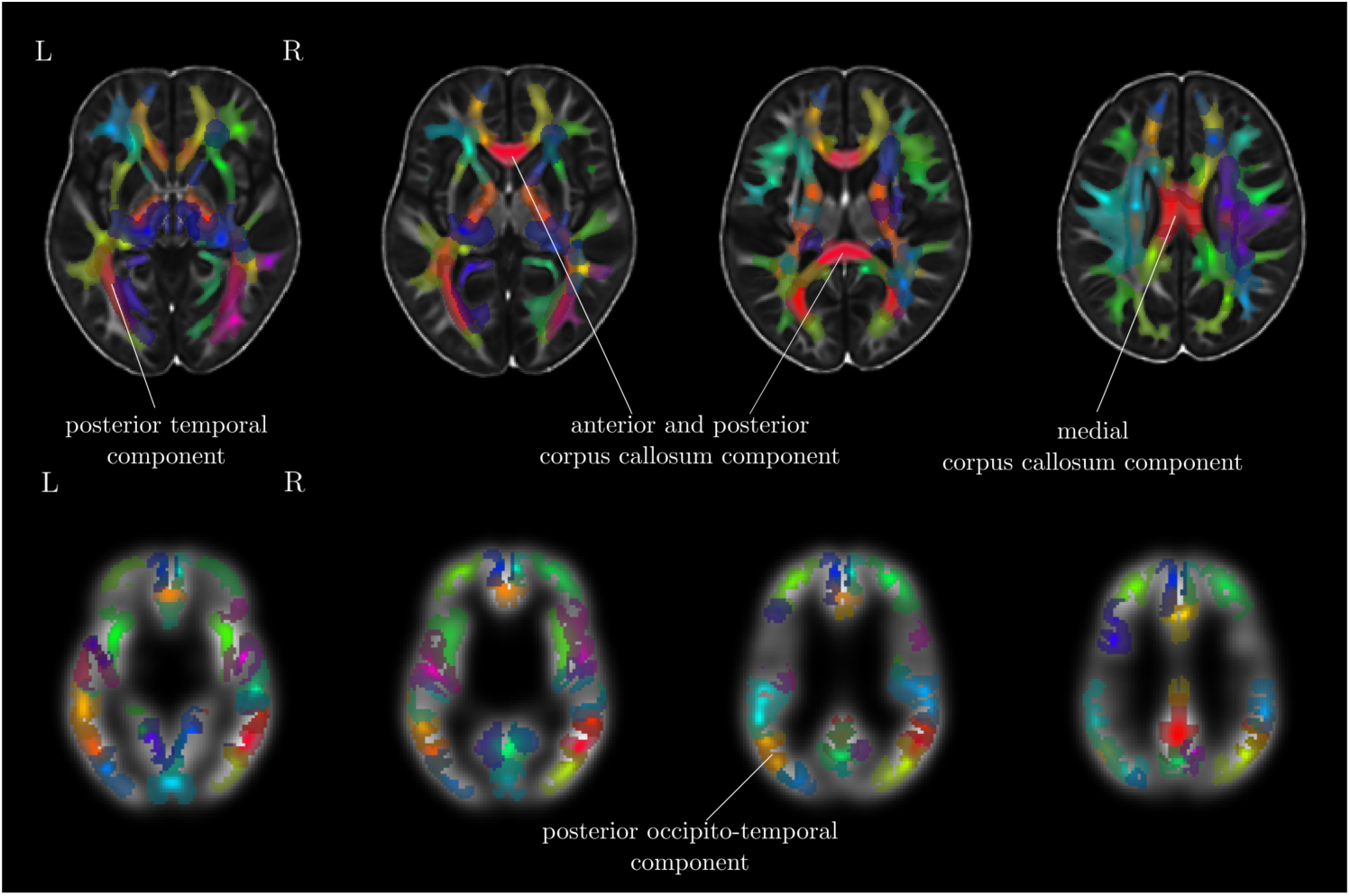
Overview of the Eigenanatomy decomposition for FA images (top) and cortical thickness maps (bottom). The 32 components indicated by eigenatomy decompositon are shown on top of the study-specific FA and cortical thickness template. Cortical thickness images were downsampled and smoothed. Labels indicate the components that were found to show interactions with working memory scores and age.

### 3.7 Statistical analysis

We wanted to test how brain morphology was associated with the components of the working memory system, and the extent to which this relationship was moderated by age. The relationship between these factors was therefore tested in the following set of regression models: a) age predicting working memory performance, b) age predicting brain morphology measures, c) brain morphology predicting working memory; and ultimately d) the interaction between brain morphology and age predicting working memory (see Figure 3 for an overview of these models). Gender and an intercept term were included as additional regressors in each model. Assessment of Cook’s distance (Cook, 1977) indicated no particularly influential data points in the regression models. Therefore, all available data points were retained in the analysis. Regression analysis was carried out using the ‘stats’ package v3.1.2 in Rbase.

**Figure 3:**
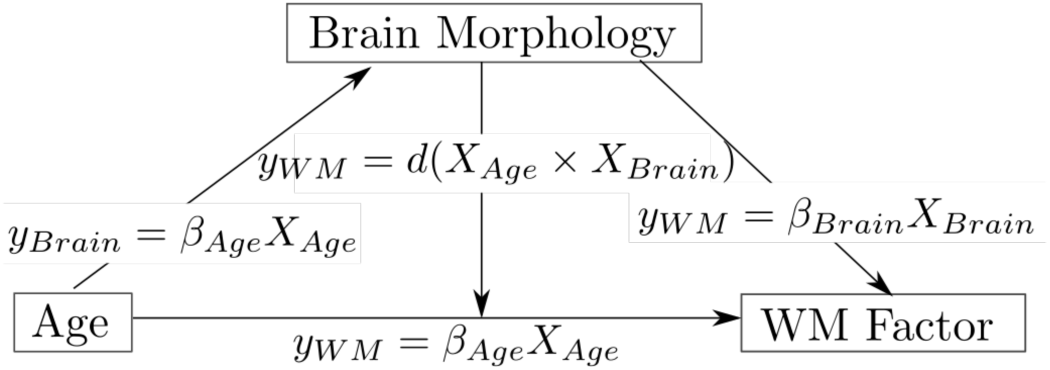
Relationships between age, brain morphology, and working memory factors explored in the current analysis. The relationship between age and working memory factors (Verbal, Executive, Spatial), age and brain morphology measures (FA, cortical thickness), and the interaction effect between age and brain morphology on working memory factors was investigated. All models further contained gender as a regressor of no interest as well as an intercept term and error term. The interaction model also contain terms for age and brain morphology separately. Models for cortical thickness analysis also contained intracranial volume as a regressor of no interest.

## 4 Results

### 4.1 Factor analysis of behavioural data

Principal component analysis (PCA) was applied to the raw scores of the working memory battery to derive the latent variable structure thought to underly working memory (A. D. Baddeley & Hitch, 1974). Assessment of Mahalanobis distance did not indicate outliers in the cognitive scores (Maximum distance D^2^(4)=15.542, critical value=18.47). Correlations between raw scores were moderate to high (range: 0.39 to 0.63). A 3-factor PCA solution with varimax rotation was selected that corresponded best with theoretical models (T. P. Alloway et al., 2004, Gathercole et al., 2004, Hornung et al., 2011, A. Baddeley, 2003, 2012, Gathercole et al., 2004). This solution explained 92% of the variance in the raw scores. The factor structure suggest a verbal, a spatial, and an executive factor. Factor loadings are shown in Table **Error! Reference source not found.**.

### 4.2 Working memory performance improves with age

Linear regression indicated that age was significantly associated with increases in working memory scores (Effects of age including gender as nuisance regressor: Verbal factor: F(2,150) = 4.538, p = 0.012, R^2^ = 0.057, R^2^_Adjusted_ = 0.044, β_Age_ = 0.010, t_Age_(150) = 2.99, p = 0.003; Executive factor: F(2,150) = 6.506, *p* = 0.002, R^2^ = 0.079, R^2^_Adjusted_ = 0.068, β_Age_ = 0.003, t_Age_(150) = 3.09, *p* = 0.002; Spatial factor: F(2,150) = 16, p < 0.001, R^2^ = 0.176, R^2^_Adjusted_ = 0.165, β_Age_ = 0.018, t_Age_(150) = 5.66, *p* < 0.001, see Figure 4a). Comparison with alternative quadratic and cubic models using the Akaike Information Criterion (AIC) as a measure of parsimony (Akaike, 1974) suggested that a linear model provided the overall best account for the relationship between age and factor scores in the current data (Verbal factor: AIC_linear_ = 433.12, AIC_quadratic_ = 432.21, AIC_cubic_ = 427.54; Executive factor: AIC_linear_ = 428.46, AIC_quadratic_ = 429.52, AIC_cubic_ = 431.51; Spatial factor: AIC_linear_ = 408.91, AIC_quadratic_ = 409.94, AIC_cubic_ = 411.61).

**Figure 4:**
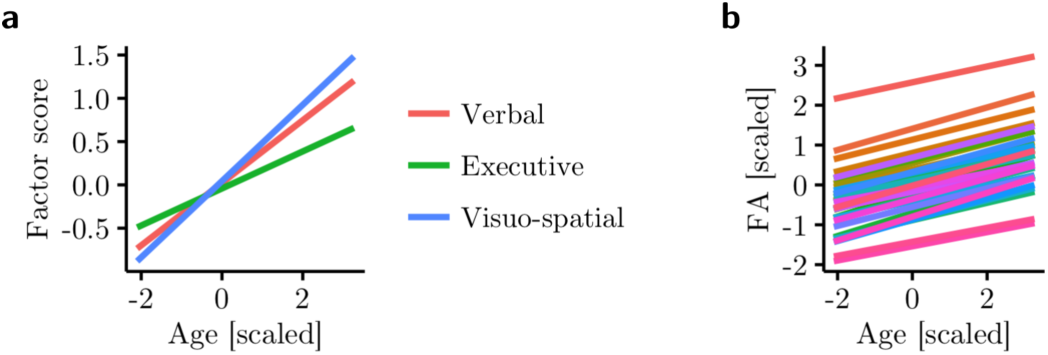
**a**: Relationship between age and verbal, executive, and visuo-spatial factor scores. Linear regression analysis indicated significantly higher scores in older participants for all factors. **b**: Relationship between age and FA within eigenanatomy components. Higher FA was significantly related to age in 30 out of 32 eigencomponents (shown).

### 4.3 FA increases with age

In order to assess the relationship between each measure of brain morphology and participant age, linear regression analysis was carried out. For FA, the effect of age was assessed including gender and an intercept in the model for all eigenanatomy components (y_FA_ = β_Age_X_Age_ + β_Gender_X_Gender_ + β_Intercept_ + ε). For FA, the results indicated a significant effect of age in 30 of the 32 components after Bonferroni correction for multiple comparisons. The effect was marginal for the remaining 2 components after correction for multiple comparisons (p < 0.051). The slopes were positive for all components (β_Age_: mean = 0.22, SD = 0.03, Range = 0.16-0.29, based on z-scores, see Figure 4b) indicating that FA increased with age for all eigenanatomy components. For cortical thickness, the model further included intracranial volume as a regressor of no interest (y_Thickness_ = β_Age_X_Age_ + β_Gender_X_Gender_ + β_ICV_X_ICV_ + β_Intercept_ + ε). The results indicated no significant relationship between age and cortical thickness. The slopes ranged from 0 to slightly positive (*beta*_Age_: mean = 0.05, SE = 0.01, Range=-0.08-0.18, based on z-scores).

### 4.4 FA predicts differences in executive scores

The relationship between brain morphology and factor scores was then assessed without taking age into account. To this end, linear regression models were fitted with the factor score as the outcome and FA and Gender as predictors on each of the assessments (y_Factor_ = β_FA_X_FA_ + β_Gender_X_Gender_ + β_Intercept_ + ε). Importantly, there was no significant effect of FA in any eigenanatomy component for the verbal and visuospatial storage factor after correction for multiple comparisons. There were significant effects of FA in 16 eigenanatomy components for the executive factor (see Table 2). For cortical thickness, the model further contained intracranial volume as a regressors of no interest (y_Factor_ = β_Thickness_X_Thickness_ + β_ICV_X_ICV_ + β_Gender_X_Gender_ + β_Intercept_ + ε). The results of the regression analysis indicated no significant effect of cortical thickness within any of the 32 eigenanatomy components on scores for any of the working memory factors (*corrected-p* > 0.05). In summary, FA predicted working memory capacity associated with the executive factor, while cortical thickness was not significantly associated with working memory performance.

**Table 1:**
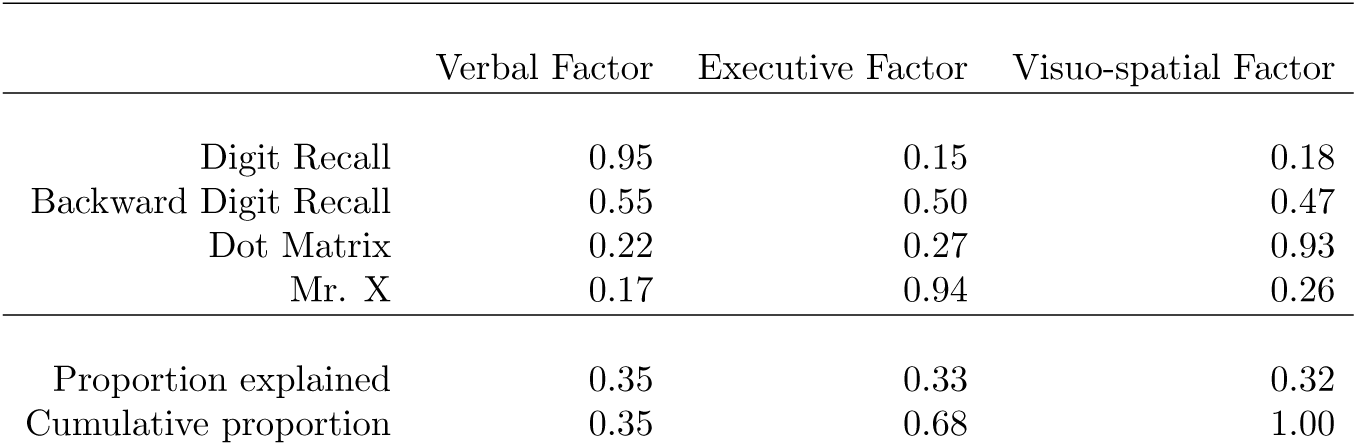
Loading of factors based on principal component analysis using varimax rotations of the raw working memory scores. The three factor solution explained 93% of the variance. The factor loadings suggested a verbal and spatial storage factor, and an executive factor.

**Table 2:**
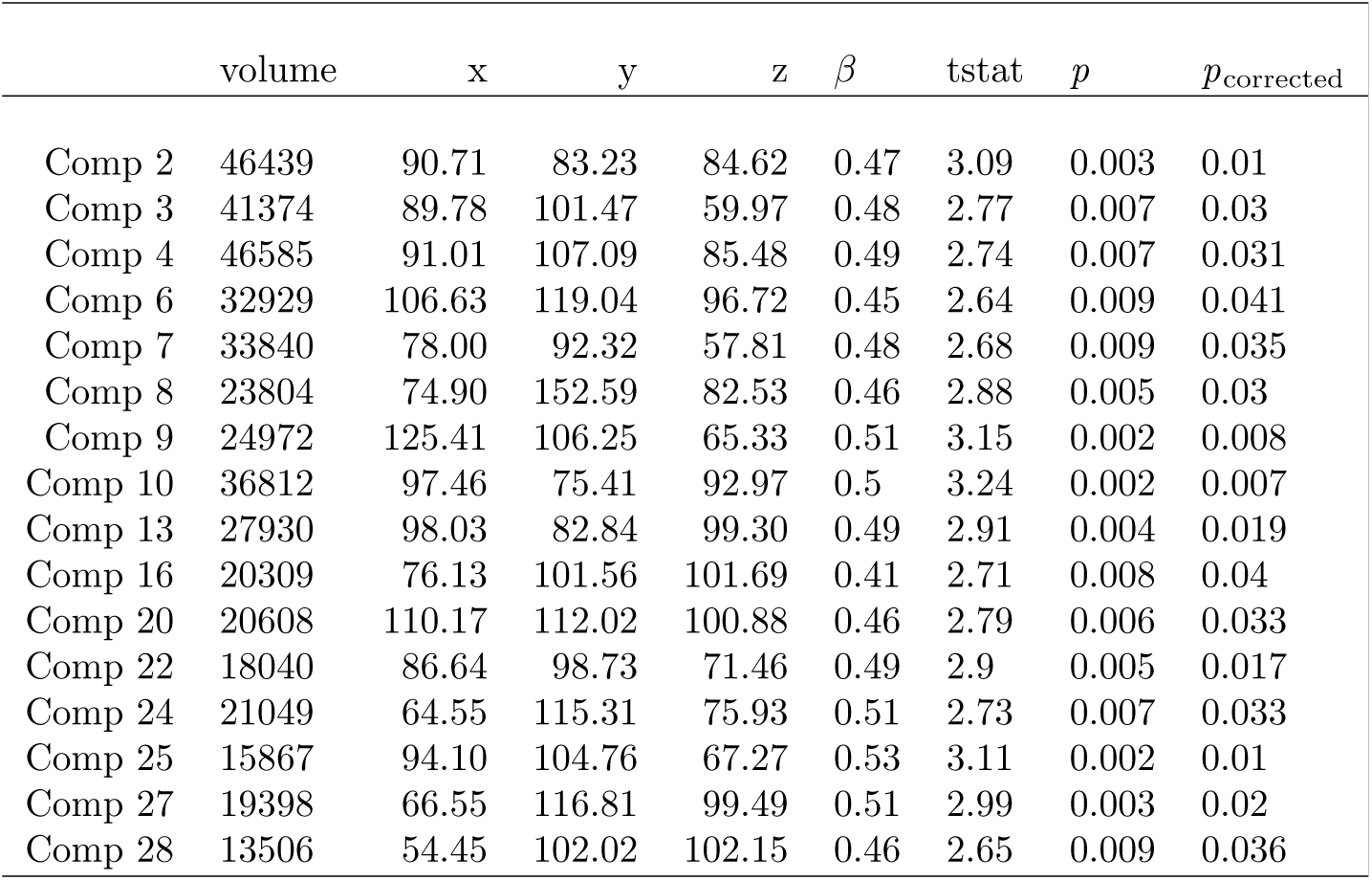
Table FA within eigenanatomy components that showed significant linear relationships with executive function scores. The coordinates refer to the position of the ROI centroid in MNI152 space.

| | volume | x | y | z | $\beta$ | tstat | $p$ | $p_{\text{corrected}}$ |
| --- | --- | --- | --- | --- | --- | --- | --- | --- |
| Comp 2 | 46439 | 90.71 | 83.23 | 84.62 | 0.47 | 3.09 | 0.003 | 0.01 |
| Comp 3 | 41374 | 89.78 | 101.47 | 59.97 | 0.48 | 2.77 | 0.007 | 0.03 |
| Comp 4 | 46585 | 91.01 | 107.09 | 85.48 | 0.49 | 2.74 | 0.007 | 0.031 |
| Comp 6 | 32929 | 106.63 | 119.04 | 96.72 | 0.45 | 2.64 | 0.009 | 0.041 |
| Comp 7 | 33840 | 78.00 | 92.32 | 57.81 | 0.48 | 2.68 | 0.009 | 0.035 |
| Comp 8 | 23804 | 74.90 | 152.59 | 82.53 | 0.46 | 2.88 | 0.005 | 0.03 |
| Comp 9 | 24972 | 125.41 | 106.25 | 65.33 | 0.51 | 3.15 | 0.002 | 0.008 |
| Comp 10 | 36812 | 97.46 | 75.41 | 92.97 | 0.5 | 3.24 | 0.002 | 0.007 |
| Comp 13 | 27930 | 98.03 | 82.84 | 99.30 | 0.49 | 2.91 | 0.004 | 0.019 |
| Comp 16 | 20309 | 76.13 | 101.56 | 101.69 | 0.41 | 2.71 | 0.008 | 0.04 |
| Comp 20 | 20608 | 110.17 | 112.02 | 100.88 | 0.46 | 2.79 | 0.006 | 0.033 |
| Comp 22 | 18040 | 86.64 | 98.73 | 71.46 | 0.49 | 2.9 | 0.005 | 0.017 |
| Comp 24 | 21049 | 64.55 | 115.31 | 75.93 | 0.51 | 2.73 | 0.007 | 0.033 |
| Comp 25 | 15867 | 94.10 | 104.76 | 67.27 | 0.53 | 3.11 | 0.002 | 0.01 |
| Comp 27 | 19398 | 66.55 | 116.81 | 99.49 | 0.51 | 2.99 | 0.003 | 0.02 |
| Comp 28 | 13506 | 54.45 | 102.02 | 102.15 | 0.46 | 2.65 | 0.009 | 0.036 |

### 4.5 The interaction between age and FA, and between age and cortical thickness predicts the executive component of WM

Finally, the relationship between brain morphology and components of the working memory system is moderated by age was investigated. The regression model for this analysis contained age, gender, FA within the eigenanatomy component, the interaction between age and component FA, and an intercept as regressors (y_Factor_ = β_Age_X_Age_ + β_FA_X_FA_ + d(X_Age_ × X_FA_) + β_Gender_X_Gender_ + β_Intercept_ + ε). The results indicated a significant effect of the interaction between age and FA on the executive factor in two eigenanatomy components (Corpus callosum component: β=−0.337, t(5)=−3.35, *p* = 0.001, *p*_corrected_=0.036; Occipitotemporal white matter component: β=−0.368, t(5)=−3.32, *p* = 0.001, *p*_corrected_=0.039, see Figure 5).

**Figure 5:**
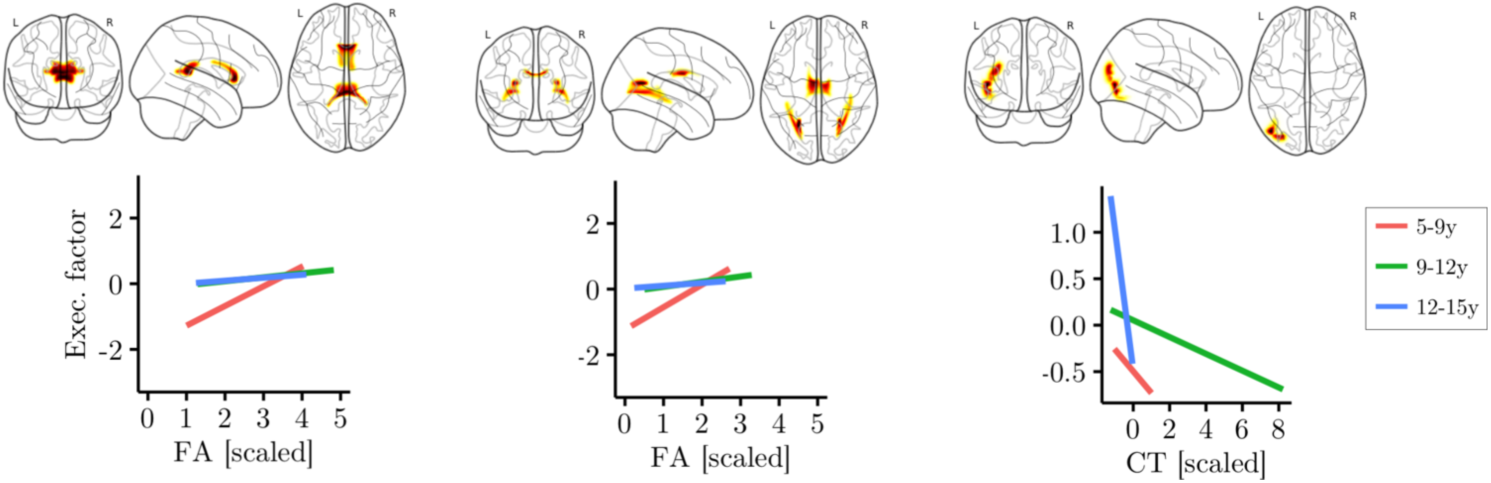
Interaction effect of age and measures of brain morphology (FA, cortical thickness) on executive factor scores. Age was split into three groups for better visualization of the results, but was treated as a continuous variable in the main analysis. Glass brain maps represent the topography of the components in MNI space. Regression analysis indicated significant interactions in two FA components (anterior and posterior corpus callosum, medial corpus callosum and bilateral posterior temporal white matter). FA in these components was more predictive of executive scores in younger children. For cortical thickness, one component in the left occipitotemporal cortex showed a significant interaction effect with age. In this component higher cortical thickness was more predictive of lower executive function scores in older children.

For cortical thickness, intracranial volume was included as an additional regressor of no interest (y_Factor_ = β_Age_X_Age_ + β_Thickness_X_Thickness_ + d(X_Age_ × X_Thickness_) + β_Gender_X_Gender_ + β_ICV_X_ICV_ + β_Intercept_ + ε). The results of the regression analysis indicated a significant interaction between age and cortical thickness for one eigenanatomy component (left temporal thickness component: β=0.56, t(5)=−0.91, *p* = 0.002, *p*_corrected_=0.049)

## 5 Discussion

The aim of the current study was to explore how structural brain correlates of components of the working memory change over developmental time. Our findings show that neural structures contributing to the executive component of the working memory system are not invariant across age, but interact with it. Specifically, the corpus callosum and bilateral posterior temporal white matter, and a cortical thickness in the left occipitotemporal cortex were found to contribute diffferently to the executive component of working memory depending on age.

### 5.1 Scores associated with the verbal, visuo-spatial, and executive factor increase with age

Rather than relying on an individual task to measure working memory performance, we used a battery of tasks to asses verbal and visuospatial short-term and working memory. We derived a factor solution that provides very good agreement with the structure expected for this particular assessment battery (T. Alloway, 2007). The loading of the tasks onto these factors was consistent with a model of working memory wherein short-term memory tasks for verbal and visuo-spatial materials load onto separate factors, and tasks with a higher executive demand load onto an additional factor (A. D. Baddeley & Hitch, 1974). Regression modelling indicated a significant linear relationship between factor scores and age for all factors. This finding is line with previous studies that indicate linear increases in short-term and working memory capacity throughout childhood and adolescence (Gathercole et al., 2004, Conklin, Luciana, Hooper, & Yarger, 2007, Swanson, 1999). Having replicated the widely reported working memory factor structure, and the age-related improvements in performance, we could then explore how changes in brain structure interact with these developmental relationships.

### 5.2 White matter organization but not cortical thickness changes with age

The second step in the analysis was to explore which aspects of neurophysiology change show the greatest degrees of age-related change. FA was significantly related to age in almost all components. Previous studies also found a positive relationship between age and FA in the mid-childhood to adolescence range for most white matter tracts (Muftuler et al., 2012, Barnea-Goraly, 2005, Qiu, Tan, Zhou, & Khong, 2008). Increasing FA is likely to reflect the contribution of different biological processes (Alexander et al., 2011), including differences in fibre organisation (Dubois et al., 2007), and increasing myelination (Dean et al., 2014, Giorgio et al., 2008).

No association between age and cortical thickness was found. This was unexpected as a number of studies reported decreasing cortical thickness with age (Wierenga, Langen, Oranje, & Durston, 2014, Sowell, 2004, Tamnes et al., 2009, Wierenga et al., 2014). However, these studies included participants from early childhood to adulthood (Tamnes et al., 2009, Sowell et al., 2006, Wierenga et al., 2014), or mapped changes longitudinally over a shorter period (Sowell, 2004, Shaw et al., 2006). The narrower age range in the current study could account for the absence of age-related changes in cortical thickness in the current study.

In summary, analysis of the relationship between age and brain morphology suggests that FA is a good indicator of brain development in 6 to 16 year age range. This suggests that the neural changes across this age span are primarily driven by the development of structural connections and integration within brain systems.

### 5.3 Brain morphology and age interact in the development of the executive component of working memory

The main analysis investigated the relationship between working memory factor scores and brain morphology and the extent to which these relationships are moderated by age. Significant interactions between age and brain morphology measures were found for the executive aspect of working memory performance in two white matter and one cortical thickness component. The white matter comprised the corpus callosum and bilateral posterior white matter, while the cortical thickness component was located in the left posterior temporal lobe. The anatomical structures identified in the current study have been previously implicated in working memory. For instance, diffusion parameters of corpus callosum subregions were found to relate to working memory capacity in children with traumatic brain injury (Treble et al., 2013) and have been reported to show changes following working memory training (Takeuchi et al., 2010). Posterior temporal white matter has also been shown to relate to working memory performance (Golestani et al., 2014, Burzynska et al., 2011).

Previous behavioural studies indicate that strategies in working memory tasks changed over development (Hitch et al., 1991, Gathercole et al., 1994). Further, the neural correlates of cognitive development show a general shift from using general resources in younger children to adult-like recruitment of specialised networks of regions in older participants (Johnson, 2011). Functional neuroimaging studies indicate that this developmental progression is also observed for working memory (Ciesielski et al., 2006, Vogan et al., 2016, Scherf, Sweeney, & Luna, 2006, Crone, Wendelken, Donohue, Leijenhorst, & Bunge, 2006). Over development, different cognitive resources and neural systems may therefore support task performance. The findings of the current study indicate that younger children benefited more from microstructural integrity of the corpus callosum and posterior temporal white matter, whereas the relationship was attentuated in older children and adolescents. In contrast, higher cortical thickness in a posterior temporal area was associated with lower levels of performance in older but not in younger children. This may suggest that the executive factor in working memory is supported by different brain systems depending on age. For younger children, communication mediated by the corpus callosum and posterior white matter is critical, while the structure of the left posterior temporal cortex is important for performance in older children.

Empirical studies grounded in the interactive specialisation view of brain development have indicated that core areas of processing are involved across the life span, but additional non-specific areas contribute in younger children. This developmental tendency has been convincingly demonstrated for face (Kadosh, Kadosh, Dick, & Johnson, 2010, Kadosh, Johnson, Henson, Dick, & Blakemore, 2013) and language processing (Weiss-Croft & Baldeweg, 2015), but working memory has not been investigated in this way so far. The current data show greater importance of white matter in corpus callosum and posterior temporal white matter in younger children for the executive component of working memory, and greater importance of left occipitotemporal cortical thickness in older children. The importance of callosal connections may indicate a reliance on a more distributed system, potentially involving recruitment of functionally homologous areas in both hemispheres, in younger children. In turn, the involvement of the left temporal lobe may indicate increased specialisation and lateralisation of the working memory system in older children. Although these final suggestions remain speculative, and to be tested.

## 6 Conclusion

The aim of the current study was to investigate how the relationship between individual differences in brain structure and working memory performance changes with age. The results indicated higher contribution of callosal and temporal white matter in younger children and higher contribution of left temporal cortex in older children for performance associated with the executive component of working memory. The current study indicates that the contribution of particular anatomical structures may change over development, potentially reflecting a shift from early reliance on a distributed system supported by long-range connections to later reliance on specialised local circuitry. The study underscores the importance of considering the crucial role that development may play in understanding brain-cognition relationships.

## References

Akaike, H. (1974). A new look at the statistical model identification. IEEE Transactions on Automatic Control, 19(6), 716–723.

Alexander, A. L., Hurley, S. A., Samsonov, A. A., Adluru, N., Hosseinbor, A. P., Mossahebi, P.,. (2011). Characterization of Cerebral White Matter Properties Using Quantitative Magnetic Resonance Imaging Stains. Brain Connectivity, 1(6), 423–446. doi:10.1089/brain.2011.0071.

Alloway, T. (2007). Automatic Working Memory Assessment.

Alloway, T. P., Gathercole, S. E., Kirkwood, H., & Elliott, J. (2008). Evaluating the validity of the Automated Working Memory Assessment. Educational Psychology, 28(7), 725–734. doi:10.1080/01443410802243828

Alloway, T. P., Gathercole, S. E., Kirkwood, H., & Elliott, J. (2009). The Cognitive and Behavioral Characteristics of Children With Low Working Memory. Child Development, 80(2), 606–621. doi: 10.1111/j.1467-8624.2009.01282.x

Alloway, T. P., Gathercole, S. E., Willis, C., & Adams, A.-M. (2004). A structural analysis of working memory and related cognitive skills in young children. Journal of Experimental Child Psychology, 87(2), 85–106. doi:10.1016/j.jecp.2003.10.002

Archibald, L. M., & Gathercole, S. E. (2006). Short-term and working memory in specific language impairment. International Journal of Language & Communication Disorders, 41(6), 675–693. doi: 10.1080/13682820500442602

Avants, B., Epstein, C., Grossman, M., & Gee, J. (2008). Symmetric diffeomorphic image registration with cross-correlation: Evaluating automated labeling of elderly and neurodegenerative brain. Medical Image Analysis, 12(1), 26–41. doi:10.1016/j.media.2007.06.004

Avants, B. B., Tustison, N. J., Stauffer, M., Song, G., Wu, B., & Gee, J. C. (2014). The Insight ToolKit image registration framework. Front. Neuroinform., 8. doi: 10.3389/fninf.2014.00044

Baddeley, A. (1987). Working Memory. New York: Oxford University Press.

Baddeley, A. (1992). Working memory. Science, 255(5044), 556–559.

Baddeley, A. (1996). Exploring the Central Executive. The Quarterly Journal of Experimental Psychology A, 49(1), 5–28. doi: 10.1080/713755608

Baddeley, A. (2003). Working memory: looking back and looking forward. Nature Reviews Neuroscience, 4(10), 829–839. doi: 10.1038/nrn1201

Baddeley, A. (2012). Working Memory: Theories Models, and Controversies. Annual Review of Psychology, 63(1), 1–29. doi: 10.1146/annurev-psych-120710-100422

Baddeley, A. D., & Hitch, G. (1974). Working Memory. In Psychology of learning and motivation (pp. 47–89). Elsevier BV.

Baddeley, A. D., & Lieberman, K. (1980). Spatial working memory. In R. S. Nickerson (Ed.), Attention and performance viii (p. 521–539). London and New York: Routledge - Taylor & Francis Group.

Barnea-Goraly, N. (2005). White Matter Development During Childhood and Adolescence: A Cross-sectional Diffusion Tensor Imaging Study. Cerebral Cortex, 15(12), 1848–1854. doi: 10.1093/cercor/bhi062

Barrouillet, P., Bernardin, S., & Camos, V. (2004). Time Constraints and Resource Sharing in Adults Working Memory Spans. Journal of Experimental Psychology: General, 133(1), 83–100. doi: 10.1037/0096-3445.133.1.83

Barrouillet, P., Gavens, N., Vergauwe, E., Gaillard, V., & Camos, V. (2009). Working memory span development: A time-based resource-sharing model account. Developmental Psychology, 45(2), 477–490. doi: 10.1037/a0014615

Bava, S., Thayer, R., Jacobus, J., Ward, M., Jernigan, T. L., & Tapert, S. F. (2010). Longitudinal characterization of white matter maturation during adolescence. Brain Research, 1327, 38–46. doi: 10.1016/j.brainres.2010.02.066

Bayliss, D. M., Jarrold, C., Gunn, D. M., & Baddeley, A. D. (2003). The complexities of complex span: explaining individual differences in working memory in children and adults. Journal of Experimental Psychology: General, 132(1), 71. doi: 10.1037/0096-3445.132.1.71

Behrens, T., Woolrich, M., Jenkinson, M., Johansen-Berg, H., Nunes, R., Clare, S.,. (2003). Characterization and propagation of uncertainty in diffusion-weighted MR imaging. Magnetic Resonance in Medicine, 50(5), 1077–1088. doi: 10.1002/mrm.10609

Benjamini, Y., & Hochberg, Y. (1995). Controlling the false discovery rate: a practical and powerful approach to multiple testing. Journal of the Royal Statistical Society, Series B, 57(1), 289–300.

Bosch, G. E. van den, Marroun, H. E., Schmidt, M. N., Tibboel, D., Manoach, D. S., Calhoun, V. D.,. (2012). Brain connectivity during verbal working memory in children and adolescents. Human Brain Mapping, 35(2), 698–711. doi: 10.1002/hbm.22193

Burgess, N., & Hitch, G. J. (1996). A connectionist model of STM for serial order in *Models of short-term memory*, 51–72.

Burzynska, A. Z., Nagel, I. E., Preuschhof, C., Li, S.-C., Lindenberger, U., Backman, L.,. (2011). Microstructure of Frontoparietal Connections Predicts Cortical Responsivity and Working Memory Performance. Cerebral Cortex, 21(10), 2261–2271. doi: 10.1093/cercor/bhq293

Cain, K., Oakhill, J., & Bryant, P. (2004). Childrens Reading Comprehension Ability: Concurrent Prediction by Working Memory Verbal Ability, and Component Skills. Journal of Educational Psychology, 96(1), 31– 42. doi: 10.1037/0022-0663.96.1.31

Camos, V., & Barrouillet, P. (2011). Developmental change in working memory strategies: From passive maintenance to active refreshing. Developmental Psychology, 47(3), 898–904. doi: 10.1037/a0023193

Carcello, J. V., & Nagy, A. L. (2004). Client size auditor specialization and fraudulent financial reporting. Managerial Auditing Journal, 19(5), 651–668. doi: 10.1108/02686900410537775

Case, R., Kurland, D., & Goldberg, J. (1982). Operational efficiency and the growth of short-term memory span. Journal of Experimental Child Psychology, 33(3), 386–404. doi: 10.1016/0022-0965(82)90054-6

Ciesielski, K. T., Lesnik, P. G., Savoy, R. L., Grant, E. P., & Ahlfors, S. P. (2006). Developmental neural networks in children performing a Categorical N-Back Task. NeuroImage, 33(3), 980–990. doi: 10.1016/j.neuroimage.2006.07.028

Clair-Thompson, H. L. S., & Gathercole, S. E. (2006). Executive functions and achievements in school: Shifting updating, inhibition, and working memory. The Quarterly Journal of Experimental Psychology, 59(4), 745–759. doi: 10.1080/17470210500162854

Colom, R., Jung, R. E., & Haier, R. J. (2007). General intelligence and memory span: Evidence for a common neuroanatomic framework. Cognitive Neuropsychology, 24(8), 867–878. doi: 10.1080/02643290701781557

Conklin, H. M., Luciana, M., Hooper, C. J., & Yarger, R. S. (2007). Working memory performance in typically developing children and adolescents: behavioral evidence of protracted frontal lobe development. Developmental Neuropsychology, 31(1), 103–128. doi: 10.1080/87565640709336889

Conway, A. R., Cowan, N., Bunting, M. F., Therriault, D. J., & Minkoff, S. R. (2002). A latent variable analysis of working memory capacity short-term memory capacity, processing speed, and general fluid intelligence. Intelligence, 30(2), 163–183. doi: 10.1016/S0160-2896(01)00096-4

Cook, R. D. (1977). Detection of Influential Observation in Linear Regression. Technometrics, 19(1), 15. doi: 10.1080/00401706.1977.10489493

Coupe, P., Yger, P., Prima, S., Hellier, P., Kervrann, C., & Barillot, C. (2008). An Optimized Blockwise Nonlocal Means Denoising Filter for 3-D Magnetic Resonance Images. IEEE Transactions on Medical Imaging, 27(4), 425–441. doi: 10.1109/TMI.2007.906087

Cowan, N. (1988). Evolving conceptions of memory storage, selective attention, and their mutual constraints within the human information processing system. Psychological Bulletin, 104, 163–191. doi: 10.1037/0033-2909.104.2.163

Cowan, N. (1999). An embedded-processes model of working memory. In A. Miyake & P. Shah (Eds.), Models of working memory: mechanisms of active maintenance and executive control (p. 62–101). Cambridge, UK: Cambridge University Press.

Cowan, N. (2013). Working Memory Underpins Cognitive Development Learning, and Education. Educational Psychology Review, 26(2), 197– 223. doi: 10.1007/s10648-013-9246-y

Cowan, N. (2016). Working Memory Maturation: Can We Get at the Essence of Cognitive Growth? Perspectives on Psychological Science, 11(2), 239–264. doi: 10.1177/1745691615621279

Cowan, N., Morey, C. C., AuBuchon, A. M., Zwilling, C. E., & Gilchrist, A. L. (2010). Seven-year-olds allocate attention like adults unless working memory is overloaded. Developmental Science, 13(1), 120–133. doi: 10.1111/j.1467-7687.2009.00864.x

Cowan, N., Ricker, T. J., Clark, K. M., Hinrichs, G. A., & Glass, B. A. (2014). Knowledge cannot explain the developmental growth of working memory capacity. Developmental Science, 18(1), 132–145. doi: 10.1111/desc.12197

Crone, E. A., Wendelken, C., Donohue, S., Leijenhorst, L. van, & Bunge, S. A. (2006). Neurocognitive development of the ability to manipulate information in working memory. Proceedings of the National Academy of Sciences, 103(24), 9315–9320. doi: 10.1073/pnas.0510088103

Dark, V. J., & Benbow, C. P. (1994). Type of stimulus mediates the relationship between working-memory performance and type of precocity. Intelligence, 19(3), 337–357. doi: 10.1016/0160-2896(94)90006-X

Das, S. R., Avants, B. B., Grossman, M., & Gee, J. C. (2009). Registration based cortical thickness measurement. NeuroImage, 45(3), 867–879.

Dean, D. C., O’Muircheartaigh, J., Dirks, H., Waskiewicz, N., Walker, L., Doernberg, E.,. (2014). Characterizing longitudinal white matter development during early childhood. Brain Structure and Function, 220(4), 1921–1933. doi: 10.1016/j.neuroimage.2008.12.016

Dennis, M., Agostino, A., Roncadin, C., & Levin, H. (2009). Theory of mind depends on domain-general executive functions of working memory and cognitive inhibition in children with traumatic brain injury. Journal of Clinical and Experimental Neuropsychology, 31(7), 835–847. doi: 10.1080/13803390802572419

Dubois, J., Dehaene-Lambertz, G., Perrin, M., Mangin, J.-F., Cointepas, Y., Duchesnay, E.,. (2007). Asynchrony of the early maturation of white matter bundles in healthy infants: Quantitative landmarks revealed noninvasively by diffusion tensor imaging. Human Brain Mapping, 29(1), 14–27. doi: 10.1002/hbm.20363

Dumontheil, I., & Klingberg, T. (2011). Brain Activity during a Visuospatial Working Memory Task Predicts Arithmetical Performance 2 Years Later. Cerebral Cortex, 22(5), 1078–1085. doi: 10.1093/cercor/bhr175

Ecker, U. K., Lewandowsky, S., Oberauer, K., & Chee, A. E. (2010). The components of working memory updating: An experimental decomposition and individual differences. Journal of Experimental Psychology: Learning, Memory, and Cognition, 36(1), 170. doi: 10.1037/a0017891

Engle, R. W. (2002). Working Memory Capacity as Executive Attention. Current Directions in Psychological Science, 11(1), 19–23. doi: 10.1111/1467-8721.00160

Engle, R. W., Tuholski, S. W., Laughlin, J. E., & Conway, A. R. (1999). Working memory, short-term memory, and general fluid intelligence: a latent-variable approach. Journal of Experimental Psychology: General, 128(3), 309. doi: 10.1037/0096-3445.128.3.309

Ericsson, K. A., & Kintsch, W. (1995). Long-term working memory. Psychological Review, 102(2), 211–245.

Faridi, N., Karama, S., Burgaleta, M., White, M. T., Evans, A. C., Fonov, V. (2015). Neuroanatomical correlates of behavioral rating versus performance measures of working memory in typically developing children and adolescents. Neuropsychology, 29(1), 82–91. doi: 10.1037/neu0000079

Fischl, B., & Dale, A. M. (2000). Measuring the thickness of the human cerebral cortex from magnetic resonance images. Proceedings of the National Academy of Sciences, 97(20), 11050–11055. doi: 10.1073/pnas.200033797

Fjell, A. M., Grydeland, H., Krogsrud, S. K., Amlien, I., Rohani, D. A., Ferschmann, L. (2015). Development and aging of cortical thickness correspond to genetic organization patterns. Proceedings of the National Academy of Sciences, 112(50), 15462–15467. doi: 10.1073/pnas.1508831112

Garyfallidis, E., Brett, M., Amirbekian, B., Rokem, A., Walt, S. van der, Descoteaux, M.,. (2014). Dipy a library for the analysis of diffusion MRI data. Front. Neuroinform., 8. doi: 10.3389/fninf.2014.00008

Gathercole, S. E., Adams, A.-M., & Hitch, G. J. (1994). Do young children rehearse? An individual-differences analysis. Memory & Cognition, 22(2), 201–207. doi: 10.3758/BF03208891

Gathercole, S. E., & Baddeley, A. D. (1989). Evaluation of the role of phonological STM in the development of vocabulary in children: A longitudinal study. Journal of memory and language, 28(2), 200–213. doi: 10.1016/0749-596X(89)90044-2

Gathercole, S. E., Pickering, S. J., Ambridge, B., & Wearing, H. (2004). The Structure of Working Memory From 4 to 15 Years of Age. Developmental Psychology, 40(2), 177–190. doi: 10.1037/0012-1649.40.2.177

Gathercole, S. E., Pickering, S. J., Knight, C., & Stegmann, Z. (2003). Working memory skills and educational attainment: evidence from national curriculum assessments at 7 and 14 years of age. Applied Cognitive Psychology, 18(1), 1–16. doi: 10.1002/acp.934

Gathercole, S. E., Tiffany, C., Briscoe, J., & Thorn, A. (2005). Developmental consequences of poor phonological short-term memory function in childhood: a longitudinal study. J Child Psychol & Psychiat, 46(6), 598–611. doi: 10.1111/j.1469-7610.2004.00379.x

Giedd, J. N., & Rapoport, J. L. (2010). Structural MRI of Pediatric Brain Development: What Have We Learned and Where Are We Going? Neuron, 67(5), 728–734. doi: 10.1016/j.neuron.2010.08.040

Giorgio, A., Watkins, K., Douaud, G., James, A., James, S., Stefano, N. D.,. (2008). Changes in white matter microstructure during adolescence. NeuroImage, 39(1), 52–61. doi: 10.1016/j.neuroimage.2007.07.043

Gogtay, N., Giedd, J. N., Lusk, L., Hayashi, K. M., Greenstein, D., Vaituzis, A. C.,. (2004). Dynamic mapping of human cortical development during childhood through early adulthood. Proceedings of the National Academy of Sciences, 101(21), 8174–8179. doi: 10.1073/pnas.0402680101

Golestani, A. M., Miles, L., Babb, J., Castellanos, F. X., Malaspina, D., & Lazar, M. (2014). Constrained by Our Connections: White Matters Key Role in Interindividual Variability in Visual Working Memory Capacity. Journal of Neuroscience, 34(45), 14913–14918. doi: 10.1523/JNEUROSCI.2317-14.2014

Hitch, G. J., Halliday, M., Schaafstal, A. M., & Heffernan, T. M. (1991). Speech “inner speech,” and the development of short-term memory: Effects of picture-labeling on recall. Journal of Experimental Child Psychology, 51(2), 220–234. doi: 10.1016/0022-0965(91)90033-O

Holmes, J., Hilton, K. A., Place, M., Alloway, T. P., Elliott, J. G., & Gathercole, S. E. (2014). Children with low working memory and children with ADHD: same or different? Frontiers in Human Neuroscience, 8. doi: 10.3389/fnhum.2014.00976

Hornung, C., Brunner, M., Reuter, R. A., & Martin, R. (2011). Childrens working memory: Its structure and relationship to fluid intelligence. Intelligence, 39(4), 210–221. doi: 10.1016/j.intell.2011.03.002

Huizinga, M., Dolan, C. V., & Molen, M. W. van der. (2006). Age-related change in executive function: Developmental trends and a latent variable analysis. Neuropsychologia, 44(11), 2017–2036. doi: 10.1016/j.neuropsychologia.2006.01.010

Imperati, D., Colcombe, S., Kelly, C., Martino, A. D., Zhou, J., Castellanos, F. X.,. (2011). Differential Development of Human Brain White Matter Tracts. PLoS ONE, 6(8), e23437. doi: 10.1371/journal.pone.0023437

Johansen-Berg, H., Behrens, T. E. J., Robson, M. D., Drobnjak, I., Rushworth, M. F. S., Brady, J. M.,. (2004). Changes in connectivity profiles define functionally distinct regions in human medial frontal cortex. Proceedings of the National Academy of Sciences, 101(36), 13335–13340. doi: 10.1073/pnas.0403743101

Johnson, M. H. (2011). Interactive Specialization: A domain-general framework for human functional brain development? Developmental Cognitive Neuroscience, 1(1), 7–21. doi: 10.1016/j.dcn.2010.07.003

Kadosh, K. C., Johnson, M. H., Henson, R. N., Dick, F., & Blakemore, S.-J. (2013). Differential face-network adaptation in children adolescents and adults. NeuroImage, 69, 11–20. doi: 10.1016/j.neuroimage.2012.11.060

Kadosh, K. C., Kadosh, R. C., Dick, F., & Johnson, M. H. (2010). Developmental Changes in Effective Connectivity in the Emerging Core Face Network. Cerebral Cortex, 21(6), 1389–1394. doi: 10.1093/cercor/bhq215

Kaiser, H. F. (1958). The varimax criterion for analytic rotation in factor analysis. Psychometrika, 23(3), 187–200. doi: 10.1007/BF02289233

Kandel, B. M., Wang, D. J., Gee, J. C., & Avants, B. B. (2015). Eigenanatomy: Sparse dimensionality reduction for multi-modal medical image analysis. Methods, 73, 43–53. doi: 10.1016/j.ymeth.2014.10.016

Kane, M. J., Conway, A. R. A., Hambrick, D. Z., & Engle, R. W. (2007). Variation in working memory capacity as variation in executive attention and control. In A. R. A. Conway, C. Jarrold, M. J. Kane, A. Miyake, & J. N. Towse (Eds.), Variation in working memory (p. 21–48). New York: Oxford University Press.

Klein, A., Andersson, J., Ardekani, B. A., Ashburner, J., Avants, B., Chiang, M.-C. (2009). Evaluation of 14 nonlinear deformation algorithms applied to human brain MRI registration. NeuroImage, 46(3), 786–802. doi: 10.1016/j.neuroimage.2008.12.037

Klingberg, T. (2006). Development of a superior frontal–intraparietal network for visuo-spatial working memory. Neuropsychologia, 44(11), 2171–2177. doi: 10.1016/j.neuropsychologia.2005.11.019

Kwon, H., Reiss, A. L., & Menon, V. (2002). Neural basis of protracted developmental changes in visuo-spatial working memory. Proceedings of the National Academy of Sciences, 99(20), 13336–13341. doi: 10.1073/pnas.162486399

Lawson, G. M., Duda, J. T., Avants, B. B., Wu, J., & Farah, M. J. (2013). Associations between childrens socioeconomic status and prefrontal cortical thickness. Developmental Science, 16(5), 641–652. doi: 10.1111/desc.12096

Lee, S.-H., Kravitz, D. J., & Baker, C. I. (2013). Goal-dependent dissociation of visual and prefrontal cortices during working memory. Nature Neuroscience, 16(8), 997–999. doi: 10.1038/nn.3452

Lenroot, R. K., Schmitt, J. E., Ordaz, S. J., Wallace, G. L., Neale, M. C., Lerch, J. P. (2009). Differences in genetic and environmental influences on the human cerebral cortex associated with development during childhood and adolescence. Human Brain Mapping, 30(1), 163–174. doi: 10.1002/hbm.20494

Logie, R. H. (1986). Visuo-spatial processing in working memory. The Quarterly Journal of Experimental Psychology Section A, 38(2), 229– 247. doi: 10.1080/14640748608401596

Luciana, M., Conklin, H. M., Hooper, C. J., & Yarger, R. S. (2005). The Development of Nonverbal Working Memory and Executive Control Processes in Adolescents. Child Development, 76(3), 697–712. doi: 10.1111/j.1467-8624.2005.00872.x

Mahone, E. M., Martin, R., Kates, W. R., Hay, T., & Horska, A. (2009). Neuroimaging correlates of parent ratings of working memory in typically developing children. Journal of the International Neuropsychological Society, 15(01), 31. doi: 10.1017/S1355617708090164

Martinussen, R., Hayden, J., Hogg-Johnson, S., & Tannock, R. (2005). A Meta-Analysis of Working Memory Impairments in Children With Attention-Deficit/Hyperactivity Disorder. Journal of the American Academy of Child & Adolescent Psychiatry, 44(4), 377–384. doi: 10.1097/01.chi.0000153228.72591.73

Michael, P. (2010). Mid-Dorsolateral Prefrontal Cortex and Posterior Parietal Cortex: Anatomical Relations and Function in Working Memory. Frontiers in Human Neuroscience, 4. doi: 10.3389/conf.fnins.2010.14.00038

Montgomery, J. W. (2000). Verbal working memory and sentence comprehension in children with specific language impairment. Journal of Speech, Language, and Hearing Research, 43(2), 293–308. doi: 10.1044/jslhr.4302.293

Muftuler, L. T., Davis, E. P., Buss, C., Solodkin, A., Su, M. Y., Head, K. M.,. (2012). Development of white matter pathways in typically developing preadolescent children. Brain Research, 1466, 33–43. doi: 10.1016/j.brainres.2012.05.035

Murphy, K., Ginneken, B. van, Reinhardt, J. M., Kabus, S., Ding, K., Deng, X.,. (2011). Evaluation of Registration Methods on Thoracic CT: The EMPIRE10 Challenge. IEEE Transactions on Medical Imaging, 30(11), 1901–1920. doi: 10.1109/TMI.2011.2158349

Nagel, B. J., Herting, M. M., Maxwell, E. C., Bruno, R., & Fair, D. (2013). Hemispheric lateralization of verbal and spatial working memory during adolescence. Brain and Cognition, 82(1), 58–68. doi: 10.1016/j.bandc.2013.02.007

Oberauer, K., Lewandowsky, S., Farrell, S., Jarrold, C., & Greaves, M. (2012). Modeling working memory: an interference model of complex span. Psychonomic Bulletin & Review, 19(5), 779–819. doi: 10.3758/s13423-012-0272-4

Oberauer, K., Süß, H.-M., Schulze, R., Wilhelm, O., & Wittmann, W. (2000). Working memory capacity — facets of a cognitive ability construct. Personality and Individual Differences, 29(6), 1017–1045. doi: 10.1016/S0191-8869(99)00251-2

Østby, Y., Tamnes, C. K., Fjell, A. M., & Walhovd, K. B. (2011). Morphometry and connectivity of the fronto-parietal verbal working memory network in development. Neuropsychologia, 49(14), 3854–3862. doi:10.1016/j.neuropsychologia.2011.10.001

Owen, A. M., McMillan, K. M., Laird, A. R., & Bullmore, E. (2005). N-back working memory paradigm: A meta-analysis of normative functional neuroimaging studies. Human Brain Mapping, 25(1), 46–59. doi: 10.1002/hbm.20131

Peters, B. D., Szeszko, P. R., Radua, J., Ikuta, T., Gruner, P., DeRosse, P. (2012). White Matter Development in Adolescence: Diffusion Tensor Imaging and Meta-Analytic Results. Schizophrenia Bulletin, 38(6), 1308–1317. doi: 10.1093/schbul/sbs054

Poldrack, R. A., & Yarkoni, T. (2016). From Brain Maps to Cognitive Ontologies: Informatics and the Search for Mental Structure. Annual Review of Psychology, 67(1), 587–612. doi: 10.1146/annurev-psych-122414-033729

Qiu, D., Tan, L.-H., Zhou, K., & Khong, P.-L. (2008). Diffusion tensor imaging of normal white matter maturation from late childhood to young adulthood: Voxel-wise evaluation of mean diffusivity fractional anisotropy, radial and axial diffusivities, and correlation with reading development. NeuroImage, 41(2), 223–232. doi: 10.1016/j.neuroimage.2008.02.023

R Development Core Team. (2008). R: A Language and Environment for Statistical Computing. Vienna, Austria. (ISBN 3-900051-07-0)

Rossi, S., Lubin, A., Simon, G., Lanoë, C., Poirel, N., Cachia, A.,. (2013). Structural brain correlates of executive engagement in working memory: Childrens inter-individual differences are reflected in the anterior insular cortex. Neuropsychologia, 51(7), 1145–1150. doi: 10.1016/j.neuropsychologia.2013.03.011

Rotzer, S., Loenneker, T., Kucian, K., Martin, E., Klaver, P., & Aster, M. von. (2009). Dysfunctional neural network of spatial working memory contributes to developmental dyscalculia. Neuropsychologia, 47(13), 2859–2865. doi: 10.1016/j.neuropsychologia.2009.06.009

Scherf, K. S., Sweeney, J. A., & Luna, B. (2006). Brain Basis of Developmental Change in Visuospatial Working Memory. Journal of Cognitive Neuroscience, 18(7), 1045–1058.

Schmiedek, F., Lövdén, M., & Lindenberger, U. (2014). A task is a task is a task: putting complex span, n-back, and other working memory indicators in psychometric context. Frontiers in psychology, 5, 1475. doi: 10.3389/fpsyg.2014.01475

Shaw, P., Greenstein, D., Lerch, J., Clasen, L., Lenroot, R., Gogtay, N.,. (2006). Intellectual ability and cortical development in children and adolescents. Nature, 440(7084), 676–679. doi: 10.1038/nature04513

Shelton, J. T., Elliott, E. M., Matthews, R. A., Hill, B. D., & Gouvier, W. D. (2010). The relationships of working memory, secondary memory, and general fluid intelligence: working memory is special. Journal of Experimental Psychology: Learning, Memory, and Cognition, 36(3), 813. doi: 10.1037/a0019046

Siegel, L. S., & Ryan, E. B. (1988). Development of grammatical-sensitivity phonological, and short-term memory skills in normally achieving and learning disabled children. Developmental Psychology, 24(1), 28–37. doi: 10.1037/0012-1649.24.1.28

Smith, S. M., Jenkinson, M., Johansen-Berg, H., Rueckert, D., Nichols, T. E., Mackay, C. E.,. (2006). Tract-based spatial statistics: Voxelwise analysis of multi-subject diffusion data. NeuroImage, 31(4), 1487–1505. doi: 10.1016/j.neuroimage.2006.02.024

Smith-Spark, J. H., & Fisk, J. E. (2007). Working memory functioning in developmental dyslexia. Memory, 15(1), 34–56. doi: 10.1080/09658210601043384

Sowell, E. R. (2004). Longitudinal Mapping of Cortical Thickness and Brain Growth in Normal Children. Journal of Neuroscience, 24(38), 8223–8231. doi: 10.1523/JNEUROSCI.1798-04.2004

Sowell, E. R., Peterson, B. S., Kan, E., Woods, R. P., Yoshii, J., Bansal, R. (2006). Sex Differences in Cortical Thickness Mapped in 176 Healthy Individuals between 7 and 87 Years of Age. Cerebral Cortex, 17(7), 1550–1560. doi: 10.1093/cercor/bhl066

Sugase-Miyamoto, Y., Liu, Z., Wiener, M. C., Optican, L. M., & Richmond, B. J. (2008). Short-Term Memory Trace in Rapidly Adapting Synapses of Inferior Temporal Cortex. PLoS Computational Biology, 4(5), e1000073. doi: 10.1093/cercor/bhl066

Swanson, H. L. (1999). What develops in working memory? A life span perspective. Developmental Psychology, 35(4), 986–1000. doi: 10.1037/0012-1649.35.4.986

Szucs, D., Devine, A., Soltesz, F., Nobes, A., & Gabriel, F. (2013). Developmental dyscalculia is related to visuo-spatial memory and inhibition impairment. Cortex, 49(10), 2674–2688. doi: 10.1016/j.cortex.2013.06.007

Takeuchi, H., Sekiguchi, A., Taki, Y., Yokoyama, S., Yomogida, Y., Komuro, N. (2010). Training of Working Memory Impacts Structural Connectivity. Journal of Neuroscience, 30(9), 3297–3303. doi: 10.1523/JNEUROSCI.4611-09.2010

Tam, H., Jarrold, C., Baddeley, A. D., & Sabatos-DeVito, M. (2010). The development of memory maintenance: Children’s use of phonological rehearsal and attentional refreshment in working memory tasks. Journal of Experimental Child Psychology, 107(3), 306–324. doi: 10.1016/j.jecp.2010.05.006

Tamnes, C. K., Ostby, Y., Fjell, A. M., Westlye, L. T., Due-Tonnessen, P., & Walhovd, K. B. (2009). Brain Maturation in Adolescence and Young Adulthood: Regional Age-Related Changes in Cortical Thickness and White Matter Volume and Microstructure. Cerebral Cortex, 20(3), 534–548. doi: 10.1093/cercor/bhp118

Tamnes, C. K., Walhovd, K. B., Grydeland, H., Holland, D., Østby, Y., Dale, A. M.,. (2013). Longitudinal Working Memory Development Is Related to Structural Maturation of Frontal and Parietal Cortices. Journal of Cognitive Neuroscience, 25(10), 1611–1623. doi: 10.1162/jocn_a_00434

Thomason, M. E., Race, E., Burrows, B., Whitfield-Gabrieli, S., Glover, G. H., & Gabrieli, J. D. E. (2009). Development of Spatial and Verbal Working Memory Capacity in the Human Brain. Journal of Cognitive Neuroscience, 21(2), 316–332. doi: 10.1162/jocn.2008.21028

Treble, A., Hasan, K. M., Iftikhar, A., Stuebing, K. K., Kramer, L. A., Cox, C. S.,. (2013). Working Memory and Corpus Callosum Microstructural Integrity after Pediatric Traumatic Brain Injury: A Diffusion Tensor Tractography Study. Journal of Neurotrauma, 30(19), 1609–1619. doi: 10.1089/neu.2013.2934

Tustison, N. J., Cook, P. A., Klein, A., Song, G., Das, S. R., Duda, J. T.,. (2014). Large-scale evaluation of ANTs and FreeSurfer cortical thickness measurements. NeuroImage, 99, 166–179. doi: 10.1016/j.neuroimage.2014.05.044

Vestergaard, M., Madsen, K. S., Baaré, W. F. C., Skimminge, A., Ejersbo, L. R., Ramsøy, T. Z.,. (2011). White Matter Microstructure in Superior Longitudinal Fasciculus Associated with Spatial Working Memory Performance in Children. Journal of Cognitive Neuroscience, 23(9), 2135–2146. doi: 10.1162/jocn.2010.21592

Vogan, V., Morgan, B., Powell, T., Smith, M., & Taylor, M. (2016). The neurodevelopmental differences of increasing verbal working memory demand in children and adults. Developmental Cognitive Neuroscience, 17, 19–27. doi: 10.1016/j.dcn.2015.10.008

Bastian, von, C. C., & Oberauer, K. (2013). Distinct transfer effects of training different facets of working memory capacity. Journal of Memory and Language, 69(1), 36–58. doi: 10.1016/j.jml.2013.02.002

Wager, T. D., & Smith, E. E. (2003). Neuroimaging studies of working memory. Cognitive Affective, & Behavioral Neuroscience, 3(4), 255– 274. doi: 10.3758/CABN.3.4.255

Wagstyl, K., Ronan, L., Goodyer, I. M., & Fletcher, P. C. (2015). Cortical thickness gradients in structural hierarchies. NeuroImage, 111, 241–250. doi: 10.1016/j.neuroimage.2015.02.036

Weismer, S. E., Evans, J., & Hesketh, L. J. (1999). An examination of verbal working memory capacity in children with specific language impairment. Journal of Speech, Language, and Hearing Research, 42(5), 1249–1260. doi: 10.1044/jslhr.4205.1249

Weiss-Croft, L. J., & Baldeweg, T. (2015). Maturation of language networks in children: A systematic review of 22years of functional MRI. NeuroImage, 123, 269–281. doi: 10.1016/j.neuroimage.2015.07.046

Westlye, L. T., Walhovd, K. B., Dale, A. M., Bjornerud, A., Due-Tonnessen, P., Engvig, A. (2009). Life-Span Changes of the Human Brain White Matter: Diffusion Tensor Imaging (DTI) and Volumetry. Cerebral Cortex, 20(9), 2055–2068. doi: 10.1093/cercor/bhp280

Wierenga, L. M., Langen, M., Oranje, B., & Durston, S. (2014). Unique developmental trajectories of cortical thickness and surface area. NeuroImage, 87, 120–126. doi: 10.1016/j.neuroimage.2013.11.010

